# Evolutionary feedbacks for *Drosophila* aggression revealed through experimental evolution

**DOI:** 10.1101/2024.08.30.610518

**Authors:** Anna R. Girardeau, Grace E. Enochs, Julia B. Saltz

**Affiliations:** Biosciences, Rice University; Houston TX 77005, USA

**Keywords:** Indirect genetic effects, aggression, Drosophila melanogaster, evolutionary feedbacks, experimental evolution

## Abstract

Evolutionary feedbacks occur when evolution in one generation alters the environment experienced by subsequent generations, and are an expected result of Indirect Genetic Effects (IGEs). Hypotheses abound for the role of evolutionary feedbacks in climate change, agriculture, community dynamics, population persistence, social interactions, the genetic basis of evolution, and more; but evolutionary feedbacks have rarely been directly measured experimentally, leaving open questions about how feedbacks influence evolution. Using experimental evolution, we manipulated the social environment in which aggression was expressed and selected in fruit fly (*Drosophila melanogaster*) populations to allow or limit feedbacks. We selected for increased male-male aggression while allowing either positive, negative, or no feedbacks, alongside unselected controls. We show that populations undergoing negative feedbacks had the weakest evolutionary changes in aggression, while populations undergoing positive evolutionary feedbacks evolved supernormal aggression. Further, the underlying social dynamics evolved only in the negative feedbacks treatment. Our results demonstrate that IGE-mediated evolutionary feedbacks can alter the rate and pattern of behavioral evolution.

**Significance:** Evolutionary feedbacks occur when evolution in one generation alters the environment experienced by subsequent generations, thereby influencing subsequent evolution. While feedbacks are thought to be important, they have rarely been directly observed because they happen over many generations and alongside other confounding processes. Here, we studied fruit flies, *Drosophila melanogaster*. In each population, we selected the most aggressive males in experimental treatments expected to produce positive, negative, or no feedbacks. Because flies have short lifespans, we could watch evolution as it happened in the lab over 24 generations. As predicted, negative feedbacks slowed down behavioral change over generations, while positive feedbacks accelerated it. Fly “boxing” was especially affected by selection under positive feedbacks. Our results highlight how small decisions, such as retaliation, can have profound effects on evolution itself.

## Main Text

Evolutionary feedbacks occur when trait evolution in one generation alters evolutionary processes in subsequent generations (1–3). Traditional behavioral biology focused on how the environment affects behavior, often ignoring reciprocal effects whereby behaviors shape the environment (4). Yet, organisms typically—perhaps universally—alter their environments (5–7), e.g., by excreting waste, eating other organisms, and behaving socially. This interplay has important implications for evolution, because the environment mediates the genetic basis and fitness consequences of variation in traits. Consequently, the potential for evolutionary feedbacks between organisms and their environments has come into focus as a major, but largely unexplored, force driving phenotypic evolution (1, 8, 9), even on short timescales (1, 10). Hypotheses abound for the role of feedbacks in climate change (11, 12), community dynamics (13), population persistence (10), social interactions (14), the genetic basis of evolution (15–17), and more.

For social behaviors, the “environment” in which behaviors are expressed is formed by the behaviors of interacting conspecifics. Indirect genetic effects (IGEs) explicitly consider how genes expressed in these social partners may influence a focal individual’s behavior (18–21). Note that “indirect” refers here to the effect of one individual’s genes on another individual’s behaviors, and is distinct from other “indirect effects” in ecology (12). IGEs stand in contrast to “direct” genetic effects, DGEs, whereby an individual’s own genotype affects its own behavior. For example, guppies (*Poecilia reticulata*) are more willing to inspect a predator if they are paired with a partner who is also willing to inspect; and these behaviors are heritable (22). This example illustrates how social interactions may be determined by the genotypes of the participants. Indirect genetic effects are widespread, and have been identified for a wide range of traits and taxa (23–26).

Under indirect genetic effects, the environment itself can evolve: evolutionary changes in population-mean behavior also represent changes in the environment in which the behavior is expressed (18, 27). Indeed, many familiar evolutionary feedbacks are examples of evolutionary outcomes of indirect genetic effects for social behaviors, such as Fisherian runaway selection and arms races (8). The existence of these feedbacks highlights the challenge of predicting behavioral evolution over more than one generation (28–31). Yet, empirical demonstrations of such feedbacks are scarce (29, 32–34), likely because isolating the effects of evolutionary feedbacks – if any – is nearly impossible for wild populations (35).

Furthermore, theoretical predictions about how IGEs will govern evolution are most relevant to situations where the IGEs themselves remain relatively stable over generations (36). As with any evolutionary process, if the underlying mechanisms – here, ways in which the behaviors of social partners alter the behavior of a focal individual– themselves evolve, then these changes may alter future evolution. Previous studies have shown that the magnitude and direction of IGEs can readily evolve, in both the lab and the field (36, 37).

To directly identify the effect(s) of evolutionary feedbacks, we founded 12 replicate *Drosophila melanogaster* populations from the same wild base population and subjected them to 24 generations of selection for increased aggression, while allowing or limiting the opportunity for different evolutionary feedbacks. Experimental evolution is an important tool for understanding evolutionary processes, because putative causal factors may be directly manipulated under controlled conditions, and the resulting evolutionary trajectories can be carefully monitored (38–41). In experimental evolution studies, the goal is to manipulate the presence, type, or magnitude of a *single* evolutionary process to identify its effects on evolutionary outcomes. Such studies thus provide an important complement to field studies, in which multiple evolutionary processes may be acting simultaneously.

IGEs have been well-established for aggressive behavior in flies (42, 43) and other species (44, 45). Aggression is inherently a highly interactive trait, because animals plastically adjust their fighting behavior based on the responses of their opponents. Thus, the aggressive behavior of one individual is an environment that influences the aggressiveness of opponents. This plasticity occurs on two distinct timescales. On short timescales, i.e., during fights, aggression by one individual typically provokes retaliatory aggression by its opponent (in flies: (43, 46) across taxa: (45, 47)). Such **retaliation** represents a social environment that augments aggression: the more aggressive your opponent is, the more aggressive you are in return. Over longer timescales, i.e., across fights, flies who have previously experienced aggressive attacks show reduced aggressiveness in subsequent encounters (43, 48–51). **Prior experience effects** thus represent a social environment that reduces aggression: the more aggressive your first opponent is towards you, the *less* aggressive you will be in your next encounter.

These social dynamics -- retaliation and prior experience effects -- are predicted to produce potent and opposite evolutionary feedbacks (45). Selection for increased aggressiveness means that future generations will experience a highly-aggressive social environment—an environment in which retaliation will augment aggression even further. This process represents a positive feedback because the direct effect of selection and the indirect effects of changing the environment in which the behavior is expressed act in the same direction—i.e., to increase aggression, augmenting the expected evolutionary response.

To test this prediction, we had a **Positive Feedbacks (P) treatment.** To permit positive feedbacks, two naïve males from the same experimental population were paired each generation, for a total of 100 males in 50 pairs. Total aggression of each male was measured, and the 50 most aggressive individuals were selected (Figure 1). This treatment is expected to produce positive feedbacks because, as aggression evolves (i.e., the population responds to selection), males will face increasingly-aggressive opponents that will stimulate increased retaliatory responses, exaggerating the population’s response to selection.

**Figure 1.**
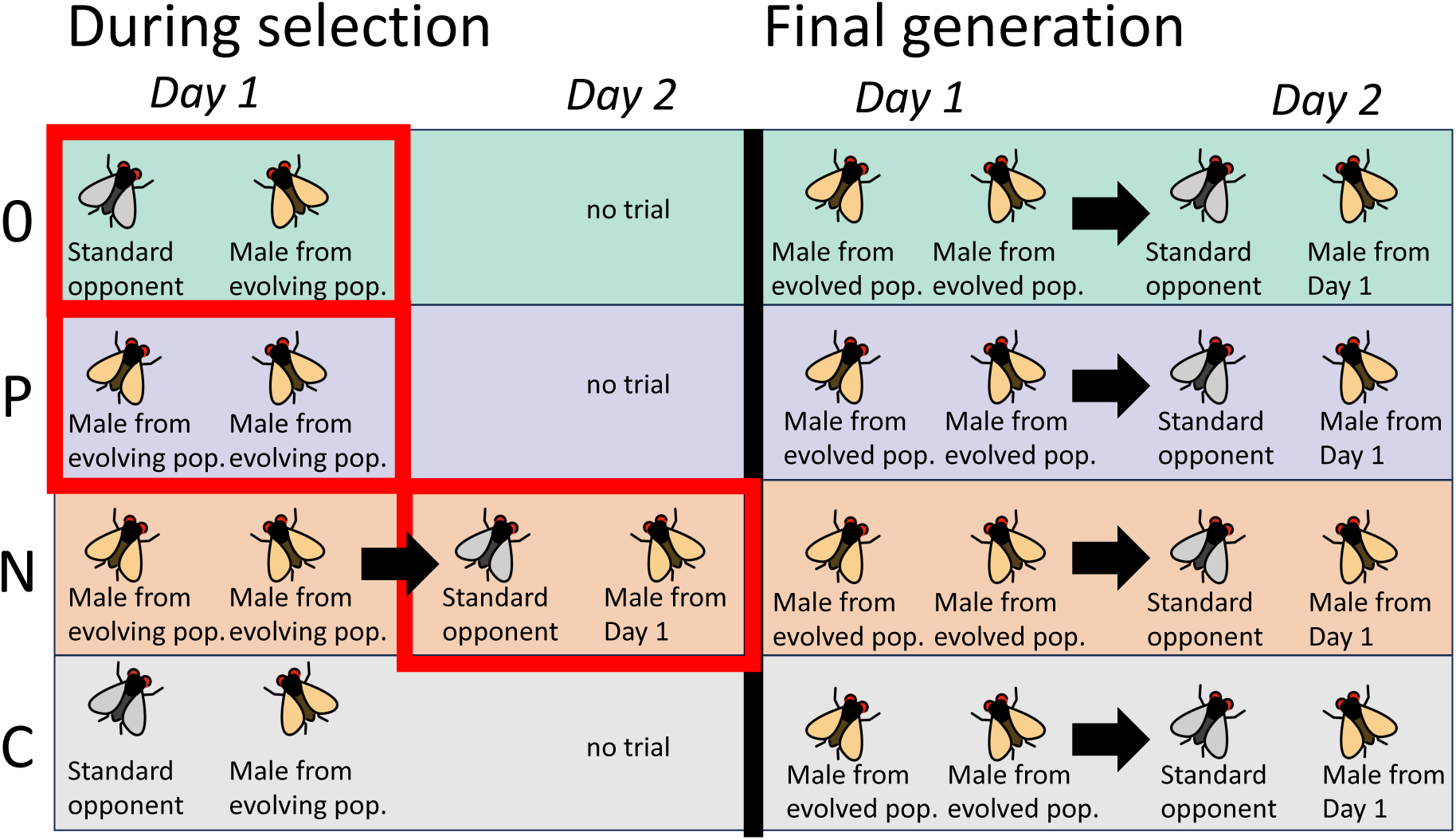
Study design and predictions. Our selection treatments manipulated the opportunity for different evolutionary feedbacks. Left: during selection (generations 1-24), 100 male flies per generation were measured for aggression in their treatment-specific social environments and times. Red boxes indicate the social environment and timing in which the 50 most aggressive males were chosen to become fathers for the next generation. Right: at the end of the experiment, all populations were measured for prior experience effects to test whether these social dynamics had themselves evolved.

In contrast, with prior experience effects, as the population-mean aggressiveness increases (i.e., the population responds to selection), more individuals will experience aggressive opponents in their first aggressive encounter, resulting in increasingly suppressed aggressiveness in subsequent encounters. This process represents a negative feedback because the direct effect of selection and the indirect effect of changing the environment in which the behavior is expressed are in opposite directions—selection acts to increase aggression, provoking more potent prior experience effects that reduce aggression, dampening the expected evolutionary response.

To test this prediction, we had a **Negative Feedbacks (N) treatment.** In this treatment, males were subjected to 2 days of behavioral analysis (Figure 1). On Day 1, two naïve males from the same experimental population were paired, for a total of 100 males in 50 pairs; this experience provides the opportunity for prior experience. The next day, Day 2, the same males were then paired with a standard opponent and aggressiveness was again measured. These standard opponents were males from the DGRP-208 inbred genotype, an inbred line whose progenitors were wild-collected and then highly inbred over many generations. Importantly, the inbred nature of these opponents mean that every male standard opponent was genetically identical to every other standard opponent in every generation. Selection was based only on males’ day 2 aggressiveness: males from the experimental population were compared for total aggression on Day 2, and the 50 most aggressive males were selected.

It may seem intuitive that “negative” feedbacks should involve selection for reduced aggression—however, this is not the case. Negative feedbacks occur when behavioral plasticity and evolutionary changes in behavior act in **opposite** directions, i.e., there is a negative covariance between behavior of opponents (18). Here, this occurs because selection for increased aggression should result in behavioral plasticity (prior experience effects) that reduces aggression.

To quantitatively identify the effects of feedbacks, we compared behavioral evolution in the **Positive feedbacks** and **Negative feedbacks** treatments to a **No feedbacks** (**0**) **treatment.** To limit opportunities for feedbacks, 100 males from the experimental population were paired with standard opponents – the same genotype of standard opponents as in the **N** treatment on day 2 (i.e., DGRP-208 inbred genotype). Males from the experimental population were compared for total aggression, and the 50 most aggressive males were selected. This treatment abrogates the opportunity for IGE-mediated evolutionary feedbacks because the standard genotype was genetically identical in every generation, meaning the social environment was not allowed to evolve.

Finally, we had **Control (C)** populations that did not undergo selection.

Each replicate was founded by collecting 4 samples of the base population, with each sample consisting of 100 males and 100 females. These 4 samples became the 4 populations from this replicate, and each one was subjected to a different experimental treatment: one population from each of the 3 selection treatments and one unselected control population. In generation 0, 100 males from each population were paired together and measured for baseline aggression to ensure similar starting points. In generations 1-24, each population was subjected to evolution for increased aggression (except unselected control populations). Each population was measured for aggression every generation as part of the selection protocol (see below), except control populations, which were measured at generation 0 and again at the midway point and at the end of the experiment. Populations within each replicate were carefully matched demographically (see Materials and Methods and Table S1) to rule out any confounding effects of population history on evolutionary change. In the final generation, samples from all populations were measured for prior experience effects so that we could test whether prior experience effects evolved (Figure 1, right), and the experiment was concluded.

By measuring 39,018 males over 25 generations in 12 initially-identical populations, our design allowed us to directly identify how evolutionary feedbacks shape changes in aggressive behavior.

## Results

In every generation, 100 males from each population (except for unselected controls) were scan-sampled (52) for aggression 24 times in one day during peak activity periods. For each observation, trained observers recorded the presence or absence of 3 behavioral components of aggression: fencing - both flies extending and engaging with each other’s legs by tapping and/or pushing; lunging - one fly rearing on his back legs and slamming his foretarsi onto his opponent; and boxing - both flies rearing on their hindlegs, “kangaroo style,” and engaging each other with their front legs by pushing and/or hitting (46). When fencing and lunging occurred, observers noted which individual was the aggressor. Boxing can only occur when both individuals are engaged. These measures are considered behavioral components of aggression (53), rather than independent behaviors, as they very often occur in a sequence during aggressive encounters (46). Finally, observers noted whether either of the flies appeared to be temporarily stuck on the food patch, as occasionally occurred.

We quantified total aggression as the unweighted sum of these behavioral components. In the selection treatments (Positive feedbacks, Negative feedbacks, and No feedbacks), the 50 males with the highest total aggression (in their treatment-specific social environment) were chosen to become fathers to the next generation alongside randomly-chosen, unmated females from their population.

After confirming that populations showed indistinguishable aggression at the beginning of the experiment (Table S3), we compared the evolutionary trajectories of total aggression across the three selection treatments and the controls using Generalized Linear Mixed Models (GLMM). Our analysis revealed a significant interaction between treatment and generation, indicating that treatments differed in how aggression changed over generations (Figure 2A, Table 1: Model 1). All 3 selection treatments evolved increased aggression, but treatments differed in the rate of behavioral change: the Negative feedbacks treatment showed slower evolutionary changes in aggression compared to the Positive feedbacks and No feedbacks treatments, while the Positive feedbacks and No feedbacks treatments showed similar rates of change, with a non-significant trend towards faster evolution in the Positive feedbacks treatment (Figure 2A, Table 2: total aggression). In contrast, unselected control populations differed significantly from all other treatments and did not evolve increased aggression (Figure 2A, Table 2: total aggression).

**Figure 2.**
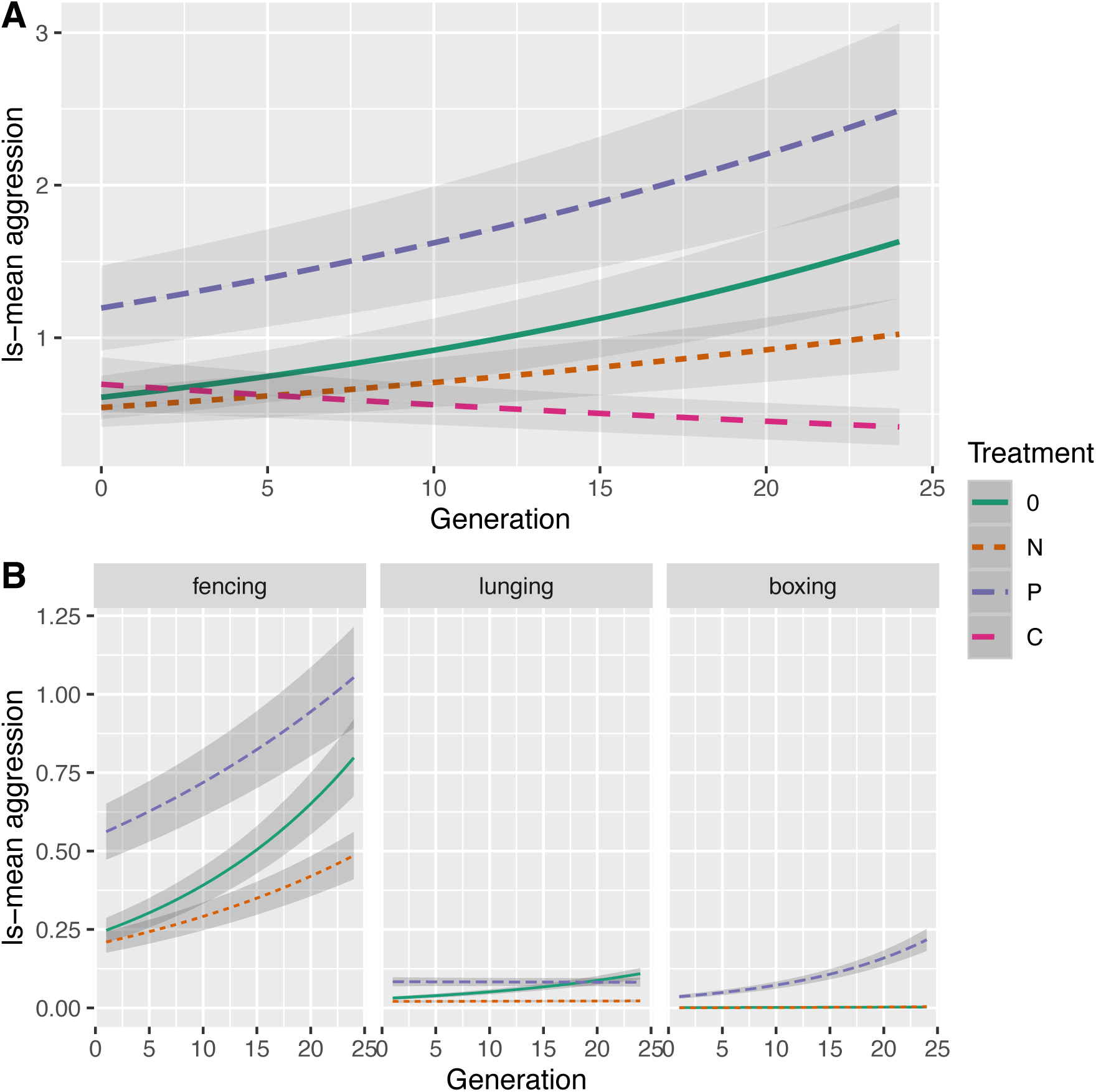
Evolutionary response to selection. **A.** Y axis is least-squares means of total aggression adjusted for other terms in the model, estimated every generation from model 1. Grey error bars indicate standard errors estimated from the model. **B.** Y axis is least-squares means for each behavioral component of aggression, adjusted for other terms in the model, estimated every generation from model 2. Note that for Boxing, the slopes for the 0 and N treatments are so low and similar that they cannot be distinguished visually on the plot. 0 indicates the No feedbacks treatment, N indicates the Negative feedbacks treatment, P indicates the Positive feedbacks treatment, and C indicates the unselected controls. N = 3 populations per treatment.

**Table 1.**
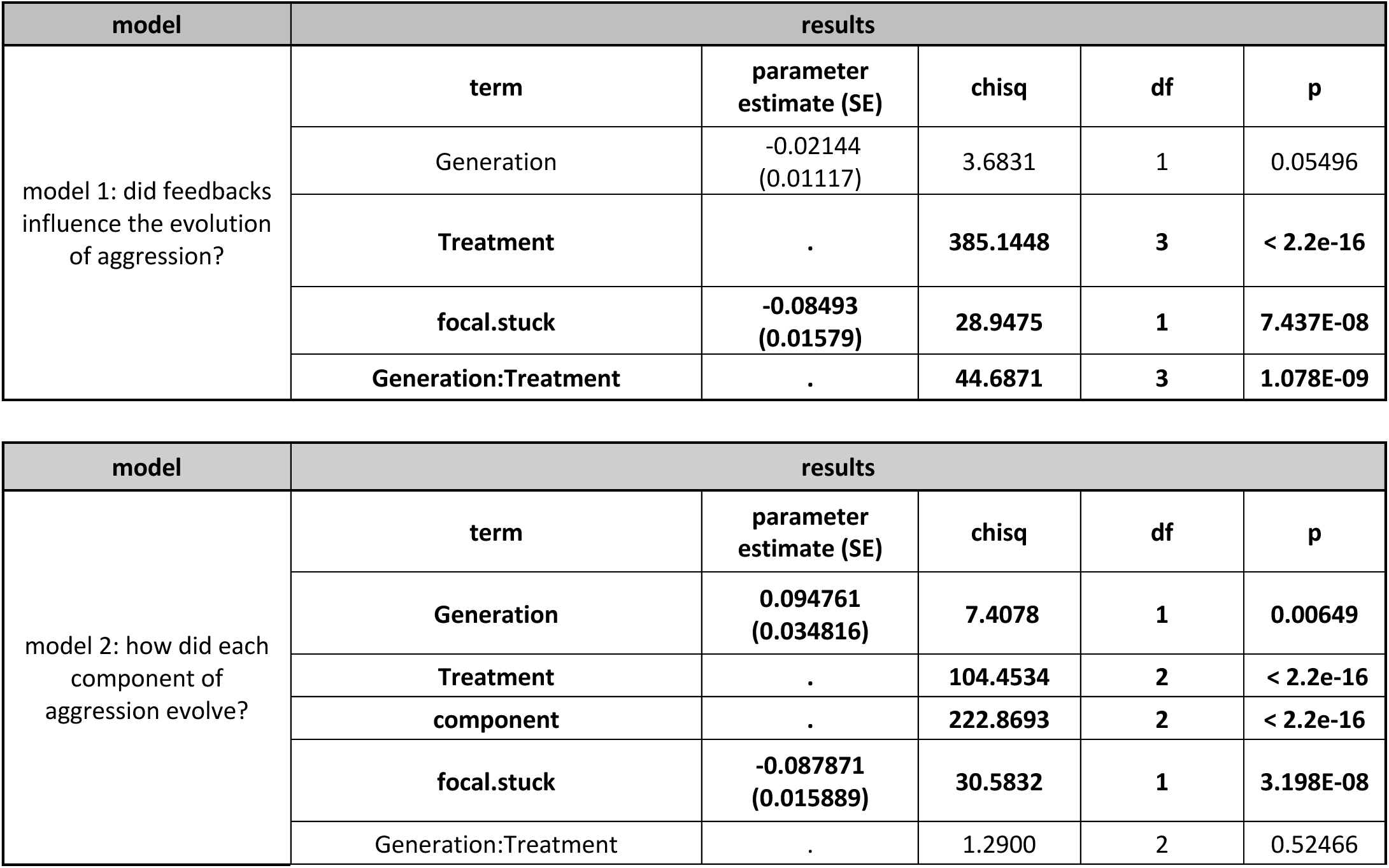
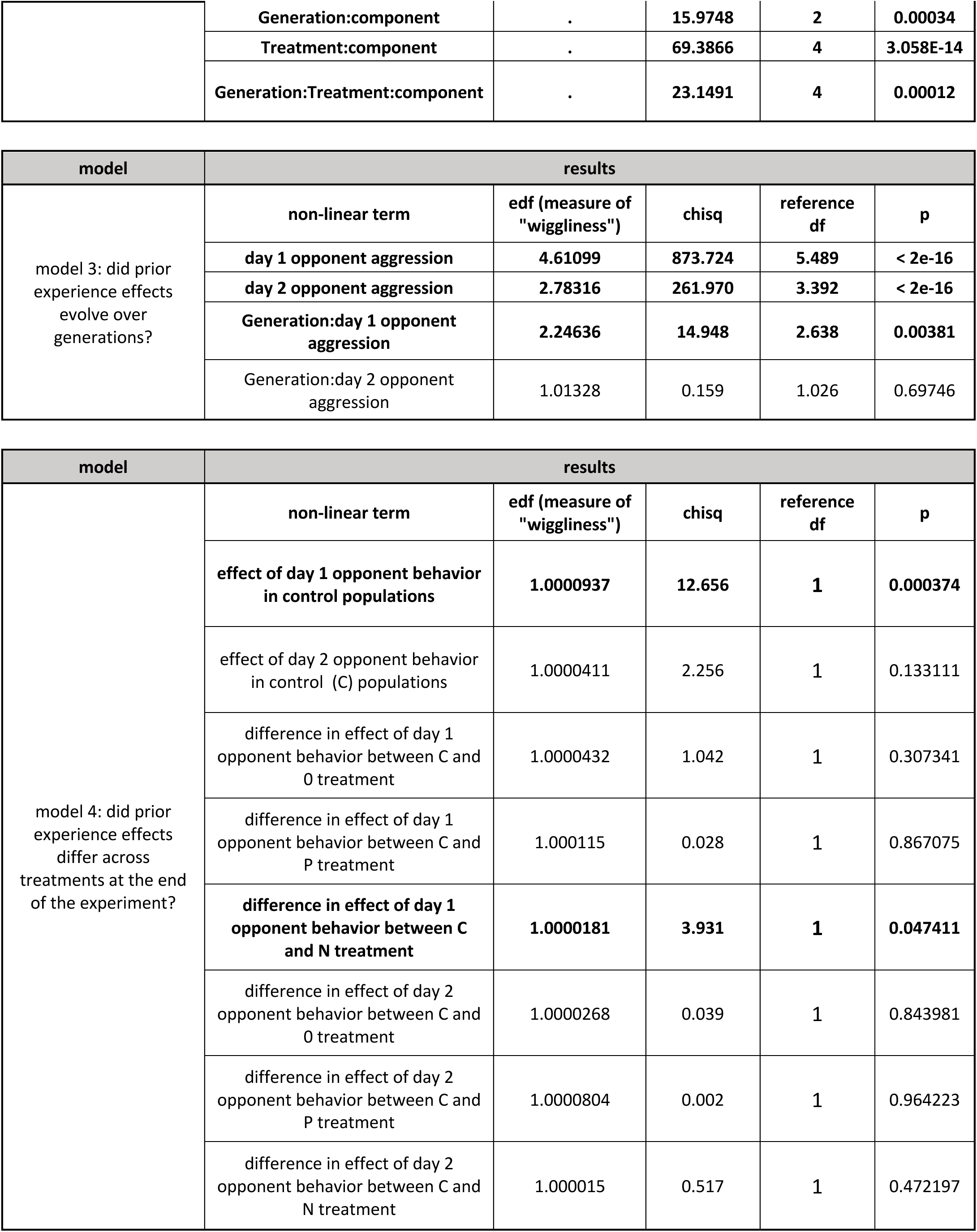
Evolutionary feedbacks influenced the evolution of aggression and prior experience effects. Important results from each model are presented. Models 1 was a GLMM describing the evolution of aggression (generations 1-24). In Model 2, we fitted a GLMM to the selected treatments only (not controls) analyzing aggression by behavioral component -- lunging, fencing, and boxing -- while accounting for repeated measurements of the same male. Model 3 was a Generalized Additive Mixed Model (GAMM) using only data from the N treatment, which is the only treatment for which prior experience effects were measured during generations 1-24. We tested whether changes in aggressive behavior over days varied across generations and due to opponent aggression on days 1 and 2 (i.e., prior experience effects). EDF describes the degree of non-linearity estimated for each term. In Model 4, we tested whether prior experience effects differed across treatments at the end of the experiment. We fitted a GAMM describing the effects of opponent aggression on each day in the Control treatment as the “reference smooth;” the other terms represent “difference smooths” that estimate differences between each treatment’s prior experience effects and those of the Control populations. For details please see Materials and Methods.

| model | results |  |  |  |  |
| --- | --- | --- | --- | --- | --- |
|  | term | parameter estimate (SE) | chisq | df | p |
| model 1: did feedbacks influence the evolution of aggression? | Generation | -0.02144<br>(0.01117) | 3.6831 | 1 | 0.05496 |
|  | Treatment | . | 385.1448 | 3 | < 2.2e-16 |
|  | focal.stuck | -0.08493<br>(0.01579) | 28.9475 | 1 | 7.437E-08 |
|  | Generation:Treatment | . | 44.6871 | 3 | 1.078E-09 |
|  | term | parameter estimate (SE) | chisq | df | p |
| model 2: how did each component of aggression evolve? | Generation | 0.094761<br>(0.034816) | 7.4078 | 1 | 0.00649 |
|  | Treatment | . | 104.4534 | 2 | < 2.2e-16 |
|  | component | . | 222.8693 | 2 | < 2.2e-16 |
|  | focal.stuck | -0.087871<br>(0.015889) | 30.5832 | 1 | 3.198E-08 |
|  | Generation:Treatment | . | 1.2900 | 2 | 0.52466 |
|  | **Generation:component** | . | **15.9748** | **2** | **0.00034** |
|  | **Treatment:component** | . | **69.3866** | **4** | **3.058E-14** |
|  | **Generation:Treatment:component** | . | **23.1491** | **4** | **0.00012** |

| **model** | **results** |  |  |  |  |
| --- | --- | --- | --- | --- | --- |
| model 3: did prior experience effects evolve over generations? | **non-linear term** | **edf (measure of "wiggliness")** | **chisq** | **reference df** | **p** |
|  | **day 1 opponent aggression** | **4.61099** | **873.724** | **5.489** | **< 2e-16** |
|  | **day 2 opponent aggression** | **2.78316** | **261.970** | **3.392** | **< 2e-16** |
|  | **Generation:day 1 opponent aggression** | **2.24636** | **14.948** | **2.638** | **0.00381** |
|  | Generation:day 2 opponent aggression | 1.01328 | 0.159 | 1.026 | 0.69746 |
| model 4: did prior experience effects differ across treatments at the end of the experiment? | **non-linear term** | **edf (measure of "wiggliness")** | **chisq** | **reference df** | **p** |
|  | **effect of day 1 opponent behavior in control populations** | **1.0000937** | **12.656** | **1** | **0.000374** |
|  | effect of day 2 opponent behavior in control (C) populations | 1.0000411 | 2.256 | 1 | 0.133111 |
|  | difference in effect of day 1 opponent behavior between C and O treatment | 1.0000432 | 1.042 | 1 | 0.307341 |
|  | difference in effect of day 1 opponent behavior between C and P treatment | 1.000115 | 0.028 | 1 | 0.867075 |
|  | **difference in effect of day 1 opponent behavior between C and N treatment** | **1.0000181** | **3.931** | **1** | **0.047411** |
|  | difference in effect of day 2 opponent behavior between C and O treatment | 1.0000268 | 0.039 | 1 | 0.843981 |
|  | difference in effect of day 2 opponent behavior between C and P treatment | 1.0000804 | 0.002 | 1 | 0.964223 |
|  | difference in effect of day 2 opponent behavior between C and N treatment | 1.000015 | 0.517 | 1 | 0.472197 |

**Table 2.** Evolutionary effects of feedbacks. Estimated marginal means of linear trends extracted from the indicated model. Slopes are presented on the scale of the original data. We tested whether each slope differed from 0 using t-tests, and we tested for slope differences between treatments. Comparisons between slopes were adjusted for multiple testing using the tukey method.

| model 1: total aggression | selection treatment | estimated trend over generations (SE) | is the slope greater than 0? | differences among treatments |
| --- | --- | --- | --- | --- |
| evolution of total aggression | C | -0.0122 (0.00663) | no (z ratio = -1.840, p = 0.0657) | a |
|  | O | 0.0422 (0.01009) | yes (z ratio = 4.183, p < 0.001) | c |
|  | N | 0.0205 (0.00536) | yes (z ratio = 3.813, p = 0.0001) | b |
|  | P | 0.0547 (0.01345) | yes (z ratio = 4.066, p < 0.0001) | c |
| model 2: behavioral components | selection treatment | estimated trend over generations (SE) | is the slope greater than 0? | differences among treatments |
| evolution of fencing | O | 2.22e-02 (3.64e-03) | yes (z ratio = 6.094, p < .0001) | a |
|  | N | 1.15e-02 (2.12e-03) | yes (z ratio = 5.425, p < .0001) | b |
|  | P | 2.08e-02 (4.11e-03) | yes (z ratio = 5.057, p < .0001) | a |
| evolution of lunging | O | 3.10e-03 (5.82e-04) | yes (z ratio = 5.333, p < .0001) | a |
|  | N | 6.73e-05 (2.07e-04) | no (z ratio = 0.325, p = 0.7453) | b |
|  | P | -6.92e-05 (4.99e-04) | no (z ratio = -0.138, p = 0.8899) | b |
| evolution of boxing | 0 | 7.49e-05 (4.95e-05) | no (z ratio = 1.513, p = 0.1303) | b |
|  | N | 1.21e-04 (1.21e-04) | yes (z ratio = 2.877, p = 0.0040) | b |
|  | P | 6.70e-03 (1.10e-03) | yes (z ratio = 6.071, p <.0001) | a |
| **model 4: prior experience effects** | **selection treatment** | **estimated effect of day 1 experience in gen 25 (SE)** | **is the slope greater than 0?** | **differences among treatments** |
| change in aggression over days | C | -0.259 (0.0727) | yes (t ratio = -3.558, p = 0.0004) | ab |
|  | 0 | -0.161 (0.0632) | yes (t ratio = -2.549, p = 0.0110) | a |
|  | N | -0.448 (0.0636) | yes (t ratio = -7.041, p <.0001) | b |
|  | P | -0.273 (0.0508) | yes (t ratio = -5.384, p <.0001) | ab |

During the experiment, it was apparent even through casual observation that the Positive feedbacks populations showed extreme levels of boxing compared to the other treatments (Figure 2B). Thus, we used a trait covariance approach to analyze how each component of aggression evolved. Lunging, fencing, and boxing were treated as repeated measures of each individual male. The resulting GLMM describing evolution in the selected populations (Model 2) revealed a significant 3-way interaction between generation, treatment, and component (Figure 2B, Table 1: Model 2), meaning that treatment differences in evolutionary change differed between fencing, lunging, and boxing. For fencing, we found that all three selection treatments evolved increased fencing; P and 0 had indistinguishable slopes, with the Negative feedbacks treatment evolving slowest (Table 2: fencing). For lunging, only the No feedbacks treatment showed significant evolutionary changes (Table 2: lunging). For boxing, we found that the Positive and Negative Feedbacks treatments showed significant evolution of boxing, but not the No feedbacks treatment (Table 2: boxing). The Positive feedbacks treatment in particular showed dramatically faster evolution of boxing compared to the other treatments; indeed, on the scale of the original data (i.e., not the latent scale), the estimated slope for the P treatment was more than 35 times greater than the corresponding estimate for the N treatment (Figure 2B, Table 2: boxing). Together, results from Models 1 and 2 demonstrate that evolutionary feedbacks alter populations’ phenotypic evolution.

Feedbacks for social behavior depend on the underlying social dynamics – here, retaliation and prior experience effects. To test whether these social dynamics changed during our experiments, we fitted additive mixed models (GAMM) that can account for non-linear effects of opponent behavior on focal male aggression. For retaliation, we tested whether the magnitude of the covariance between interacting males changed directionally over generations. While retaliation was present in all populations tested, none of the treatments showed evidence that the magnitude or form of retaliation changed over generations (Table S4).

To test for evolution of prior experience effects, we fitted a GAMM to data from the Negative feedbacks treatment (Model 3). We measured prior experience effects for each male as the change in aggression across days (i.e., focal male Day 2 aggression – focal male Day 1 aggression, (54, 55)). Our model uncovered a significant interaction between Day 1 opponent behavior and Generation (Table 1: model 3), indicating that the effects of social experience on male behavioral plasticity evolved over generations (Figure 3A – compare Generation 1 (right) to Generation 12 (center) to Generation 24 (right)). To test whether this evolutionary change in prior experience effects was unique to the Negative feedbacks treatment, we compared prior experience effects between the Positive feedbacks, Negative feedbacks, No feedbacks, and Control populations at the end of each replicate (Model 4). Our analysis (Table 1: model 4) revealed that prior experience effects differed significantly from the unselected Control populations only in the Negative feedbacks treatment (Figure 3B, Table 2: model 4).

**Figure 3.**
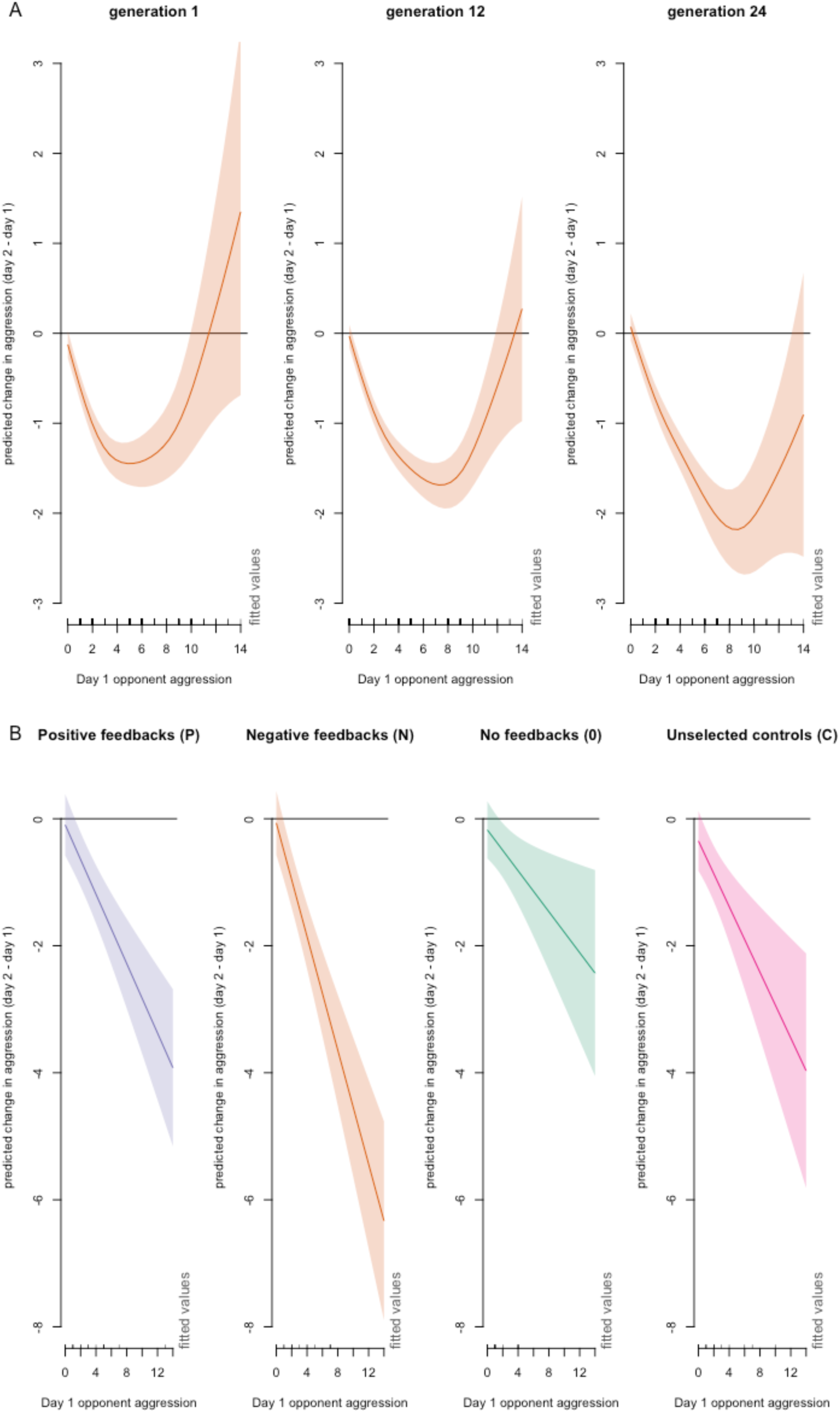
Prior experience effects evolved in the Negative Feedbacks treatment. A: prior experience effects over 24 generations in the negative feedbacks treatment. Y axis is the change in behavior over days, where positive values mean that focal male aggression was enhanced after day 1 social experience, negative values mean that focal male aggression was suppressed after day 1 social experience, and zero indicates no effect of prior experience. The x-axis is the aggression of the focal males’ day 1 opponent, and the line represents the cubic spline from Model 2 with associated standard error. Comparison of Generation 1 (A, left) with Generation 12 (A, center) and Generation 24 (A, right) illustrates that prior experience effects evolved. B: comparison of prior experience effects at the end of the experiment for each treatment. The x-axis is the aggression of the focal males’ day 1 opponent, and the line represents the cubic spline from Model 4 with associated standard error. Only the N treatment showed evolutionary changes in prior experience effects.

## Discussion

Evolutionary feedbacks are widely hypothesized to be both common and important, but quantifying the effects of evolutionary feedbacks in the midst of other population-genetic processes is challenging, leaving open questions about how evolutionary feedbacks play out in populations over time. Here, we manipulated the social environment in which aggression was expressed and selected to allow or limit the opportunity for indirect genetic effect (IGE)-mediated evolutionary feedbacks to contribute to evolution (Figure 1). We tracked the resulting evolutionary dynamics over 24 generations in 12 populations, requiring behavioral measurement of tens of thousands of males. Our results broadly support predictions (Figure 2, Tables 1-2), i.e., that positive feedbacks should accelerate phenotypic evolution while negative feedbacks should slow it (18, 19, 27, 31). As unselected control populations did not evolve increased aggression, we can be confident that these evolutionary changes are due to our experimental selection treatments. Thus, our results demonstrate that evolutionary feedbacks alter populations’ phenotypic evolution.

Not every component of aggression evolved the same way; in particular, the evolutionary changes in the Positive feedbacks treatment were dominated by the very striking evolution of supernormal boxing behavior (Figure 2B: boxing). Boxing is a mutually-expressed high-level component of aggression in which active participation by both males is required. Thus, small increases in aggressiveness over generations may have been magnified in the Positive feedbacks treatment through retaliation, making mutually-expressed, high-intensity fights substantially more common.

Evolutionary feedbacks are predicted to alter both the magnitude of selection and realized heritability (27, 31) to shape phenotypic change over time. Existing statistical approaches to disentangle these processes are focused on normally-distributed traits (56) limiting our ability to piece apart these processes in our data. We consider it most likely that IGE-mediated effects on both selection strength and realized heritability played roles in our observed results. A recent meta-analysis provides strong evidence that IGEs can alter trait heritability, particularly for behaviors, as measured through standing variation (i.e., heritability, not realized heritability, (57)). On the selection side, we performed truncation selection with the most aggressive 50% of males selected to reproduce each generation. As the populations evolved increased average aggression, the absolute number of aggressive behaviors required of a male to qualify for the selected fraction also increased. Studying evolutionary feedbacks under alternative forms of selection would be an exciting future direction.

Another key dimension of evolution under evolutionary feedbacks is that the behavioral dynamics that produce feedbacks – here, retaliation and prior experience effects – may themselves evolve. While we found no evidence for evolutionary changes in retaliation, prior experience effects showed evolutionary changes in the Negative feedbacks treatment only (Figure 3). Inspection of the reaction norms describing the effect of prior experience on later aggression, both across generations of the Negative feedbacks treatment (Figure 3A) and across treatments at the end of the experiment (Figure 3B, table 2) indicate that prior experience effects evolved to be more pronounced: being attacked by an opponent male on Day 1 produced more severe depression of aggression on Day 2, compared to generation 1 and to other treatments (Table 2, model 4). These findings suggest that negative feedbacks and their evolutionary consequences would be exacerbated under continued selection. Predicting and testing the evolution of social dynamics as indirect targets of selection is a critical future direction for understanding evolutionary feedbacks (36, 37).

## Conclusions

The existence and potency of IGE-mediated evolutionary feedbacks highlight how simple behavioral choices – such as responding to an aggressive attack – can scale up to impact the entire evolutionary process. Our results provide some of the clearest evidence to date that evolutionary changes can alter the environment to produce evolutionary feedbacks that, over time, influence evolutionary change. Together, these findings indicate that evolutionary feedbacks may not be a unique or unusual feature of evolution in specific cases, but rather, a widespread outcome of the ways that organisms interact with their environments. Thus, evolutionary feedbacks should be considered in a broad range of evolutionary studies, even when IGEs and related processes are not the explicit focus.

## Methods

### Experimental Overview

#### Replicate populations were selected for increased aggression under different social conditions, alongside unselected controls

We had 3 selection treatments representing selection for increased aggression under positive, negative, or no evolutionary feedbacks, as described below. In addition, we had control populations that did not undergo selection.

In generation 0, 100 males from each population were paired together and measured for baseline aggression to ensure similar starting points. In generations 1-24, each population was subjected to evolution for increased aggression, or not, corresponding to their assigned treatments. Each population was measured for aggression every generation as part of the selection protocol (see below), except control populations, which were measured at generation 0 and again at the midway point (generation 12 or 13). In the final generation (generation 25 for replicates 1 and 2, generation 26 for replicate 3), samples from all populations were measured for prior experience effects so that we could test whether prior experience effects evolved, and the experiment was concluded.

Replicate 1 was founded in July 2021, Replicate 2 was founded in September 2021, and Replicate 3 was founded in February 2022.

### Experimental details: flies and rearing conditions

#### Base population

As our base population, we used an outbred population of flies originally collected from Coral Gables, FL, by the de Bivort lab. Mated females were collected from the same wild population over a period of June-November 2018 and their descendants were kept at large population sizes in the de Bivort lab (61).

#### Standard opponents

We used standard opponents in several of the treatments as explained below. These were males from the DGRP-208 inbred genotype, an inbred line whose progenitors were wild-collected and then highly inbred over many generations. DGRP-208 was chosen because it showed intermediate aggression in prior studies (Hutchins et al 2024). Importantly, the inbred nature of these opponents mean that every male standard opponent was genetically identical to every other standard opponent in every generation.

Standard opponents were reared for trials by adding 10 unmated DGRP-208 females and 10 DGRP-208 males to a vial containing standard fly food. This approach minimizes variation in rearing conditions across vials. Male progeny were collected within 12 hours of eclosion and housed with other standard opponent males for 5-7 days prior to testing.

#### Rearing and preparation for aggression experiments

Experimental populations were reared on high-protein food (consisting of standard fly food supplemented with additional dead yeast to produce a 4:1 protein:carbohydrate ratio; (62)) in 6oz plastic bottles. Each bottle was founded by 25 males and 25 unmated females. The high protein food ensured that sufficiently large numbers of males eclosed around the same time, allowing us to conduct the experiment. Newly-eclosed unmated females were housed in same-sex groups. Newly-eclosed experimental males were marked with a small dot of yellow or blue acrylic paint under anesthesia so that we could distinguish individuals during behavioral trials. After painting, males were isolated for 5-7 days. This approach ensured that males were recovered from anesthesia, but had no adult social experience prior to behavioral trials.

### Selection on aggressive behavior

#### Aggression arenas

Arenas were composed of two small Petri dishes (3 cm x 1 cm) taped together. The floor of the arena was filled with standard fly food medium and a dot of active yeast. This is a typical arena design for measuring aggression in flies (e.g., (43)).

#### Measuring aggression

Because of the large number of males that needed to be measured simultaneously, we used an instantaneous scan sampling approach to measure aggression (52). Flies were added to arenas in the morning, at approximately 9:00am. Every 5 minutes during the following 60 minutes, we noted whether flies in each arena were engaged in any of 3 aggressive behaviors. A second 60-minute observation period occurred from 3:00-4:00 pm. Thus, we collected 24 total observations of each fly.

For each observation, trained observers recorded the presence or absence of 3 components of aggressive behavior: fencing - both flies extending and engaging with each other’s legs by tapping and/or pushing; lunging - one fly rearing on his back legs and slamming his foretarsi onto his opponent; and boxing - both flies rearing on their hindlegs, “kangaroo style,” and engaging each other with their front legs by pushing and/or hitting (46, 63). When lunging occurred, observers noted which individual was the aggressor. Fencing and boxing are mutually performed behaviors so both individuals were counted as engaging in each instance of these.

Finally, observers noted whether either of the flies appeared to be temporarily stuck on the food patch, as occasionally occurred. Observers did not know which flies were from which treatments while measuring aggression. To choose flies to become fathers of the next generation (in the selection treatments), we computed total aggression, which was the unweighted sum of fencing, lunging, and boxing instances observed for each individual male. For each selected population, the 50 males with the highest total aggression were selected each generation. If fewer than 50 males showed non-zero aggression, then all the males that showed any aggression were selected alongside randomly-chosen additional males until we reached 50 total males. For example, if only 45 males showed aggression, 5 males that did not show aggression (during our sampling periods) would be randomly chosen, resulting in 50 fathers for the next generation.

##### Founding the next generation

After selection, the 50 chosen males were placed into food bottles with 50 randomly chosen unmated females from their own population. The progeny of these flies became the next generation.

##### Deviations from the protocol

Due to unavoidable variability in fly eclosion rates, there were some generations in which fewer than 100 males were available from one or more of the populations. In these cases, the other populations in that replicate were subsampled to produce equal numbers of males considered for selection. For example, if one population only had 70 males available for testing, we measured those 70 males alongside 100 males from the *other* populations in that replicate. Before selection, we randomly discarded data from 30 males from the other populations, and chose the 50 most aggressive males from the remaining 70 data points.

For replicate 3 only, we had 3 generations where no selection was possible. In Generation 15, flies eclosed almost a week later than usual, possibly due to unusual weather in Houston TX. Since we were unsure of whether these flies were healthy and representative of the evolving population, we chose to allow random parents to found the next generation rather than conducting selection. Later, the standard opponent colonies became contaminated with bacteria resulting in insufficient standard opponents in generations 22, 23, and 25 (for Replicate 3 only). To ensure identical demographic histories among Replicate 3 populations, we therefore chose random flies to propagate all the populations during these “lost” generations. We concluded the experiment in Generation 26, in which we measured prior experience effects. Note that all results are adjusted for these and any other differences between Replicates.

In sum, this approach ensured identical demographic histories among populations from the same replicate. A complete record of the number of males measured for each population in each replicate is available in Table S1 in supplementary materials.

##### Final generation: test for prior experience effects

In the final generation, males from selected and control populations were reared as normal. But instead of their normal testing protocol, males from all populations were tested for prior experience effects: males from each population were paired together on Day 1 and aggression was measured, then those same males were paired with a standard opponent the next day and their aggression was measured again (i.e., as in (43, 51)).

## Analysis Methods

### Overview

We fit a series of models using data from generations 1-24 to test for evolutionary changes in aggression, retaliation, and prior experience effects, and differences in these slopes among treatments. Finally, we analyzed prior experience effect differences in the final generation of each replicate. All models included a random effect of Replicate to account for differences among replicates. A full summary of each model, the question that it answers, and its structure, is available in table S2.

In initial models, we found that the relationship between focal male aggression and opponent aggression was highly non-linear. Thus, analyses that included opponent aggression as a predictor were fit as generalized additive mixed models (GAMMs) in the package mcgv, enabling us to capture these nonlinear effects (64). Opponent aggression and its interactions were fit using thin-plate splines, which include penalties for overfitting (64). We allowed the model to fit up to 10 “knots” for each spline. Models not including the effects of opponent behavior were fit as generalized linear mixed models (GLMMs) using the package glmmTMB, enabling us to include zero-inflation terms as needed (65).

Our measure of prior experience effects focused on changes in aggression across days and was calculated as: focal male Day 2 aggression – focal male Day 1 aggression, for each male (54, 55). For these models, we used the scaled t error distribution, which is appropriate for errors that have a heavy-tailed gaussian-type distribution. For models where aggression itself was the response variable, we used a negative binomial error distribution, which is appropriate for overdispersed count data.

Preliminary models showed that fly age (i.e., whether the flies were 5, 6, or 7 days post-eclosion) was never significant, so we did not include this term in final models.

### Model fitting and inference

Residual plots from each model were examined to ensure adequate model fit. For GLMMs, residual plots were produced by simulation using the package DHARMa (66); for GAMMs, we used the gam.check function in mcgv. For GLMMs, we also used AIC analysis to decide what zero-inflation terms to include, if any.

In the final models, we used Wald type III tests (implemented in the ‘car’ package for GLMMs (67), and F tests in the “summary.gam” function in mcgv for GAMMs) to test the significance of fixed effects. We did not conduct inference on random effects.

We estimated slopes, i.e., estimated marginal means of linear trends, representing change in behavior over generations. Slopes were de-transformed to the scale of the original data, which is appropriate for interpreting evolutionary patterns (68). We tested whether each slope differed from 0 using t-tests, and we performed planned contrasts to compare slopes from different treatments. Comparisons between slopes were adjusted for multiple testing using the tukey method. These computations were implemented in the emmeans package (69).

### Model details

#### Model 1: How did aggression evolve?

To test how aggression evolved, we modeled total aggression (i.e., the unweighted sum of all aggression components) across generations. This model included data from the selection treatments as well as the controls, starting in generation 0. Fixed predictors were treatment, generation, and their interaction, with a fixed covariate indicating the number of times the fly appeared stuck on the food patch. We included terms for replicate, the date the fly was measured, the identities of the trained observers.

#### Model 2: how did each component of aggression evolve?

We used a trait covariance approach (similar to MANOVA) to analyze how total aggression and each component of aggression evolved in our experiments. For selection treatments only, we modeled “focal male aggression” as our response variable, and each individual was represented by three measurements –lunging, fencing, and boxing--with a variable, “component,” indicating which observation corresponded to which component of aggression. The model included fixed effects of generation, treatment, component, and all possible interactions among these variables. We also included a fixed covariate indicating the number of times the fly appeared stuck on the food patch. We included the same random effects as in Model 1, plus random effect of individual ID to account for the non-independence of the 3 measurements from each individual.

#### Model 3: did prior experience effects evolve in the N treatment?

In this model, we used only data from the N treatment. We tested whether within-individual changes in aggressive behavior over the 2 days of trials (measured as Day 2 aggression –Day 1 aggression for each male, see above) varied over generations and due to opponent aggression on days 1 and 2. Generation, opponent aggression on each day, and 2-way interactions between generation and opponent aggression on each day were included as fixed effects, with other terms in the model as above. In this model, an effect of a generation x opponent aggression interaction (on either day) would indicate an evolutionary change in how males responded to opponent behavior. We included the same random effects as in Model 1.

#### Model 4: did prior experience effects evolve in any of the treatments?

To compare the magnitude of prior experience effects across treatments, we tested whether changes in aggressive behavior across days (measured as Day 2 aggression – Day 1 aggression for each male, see main text) differed across treatments in Generation 25. Using GAMMs, we fitted the effects of opponent aggression on each day in the Control treatment as the “reference smooth,” and fitted effects that estimated differences between Control and each of the other treatments for these relationships as “difference smooths” (70). Significant effects of each difference smooth would indicate significant differences between this treatment’s prior experience effects and those of the control populations. We included the same random effects as in Model 1.

#### Model S1: checking assumptions

In this model we confirmed that populations had indistinguishable levels of aggression at the beginning of the experiment (generation 0) See Table S3.

#### Models S2.0, S2.N, and S2.P: did retaliation evolve?

In this series of models, we tested whether retaliation changed over generations in each selection treatment. See Table S4.

## Acknowledgments

We thank Charlotte Hovland, Autumn Hildebrand, Philip DuBose, Samitha Nemirajaiah, Vivian Ha, Jose Ramirez, Maggie Bao, Priya Trakru, Daniela Vodo, and Brooke Ermias, for logistical assistance. We thank the de Bivort lab, especially Jamilla Akhund-Zade, for supplying the outbred population.

## Author contributions

Conceptualization: JBS

Methodology: ARG, GEE, JBS

Investigation: ARG, GEE, JBS

Visualization: JBS

Funding acquisition: JBS

Project administration: ARG, GEE, JBS

Supervision: ARG, GEE, JBS

Writing – original draft: JBS

Writing – review & editing: ARG, GEE, JBS

## Reproducible research

Data and code needed to reproduce the analyses presented are included as supplemental files.

## Supplementary Materials

Supplemental Tables S1 to S4

Data files

Readme for data files

R Code

### SUPPLEMENTAL TABLES

**Table S1.** Number of males measured for aggression every generation for every population. In cases where less than 100 were measured, we matched the demographic histories of all populations in that replicate. See Materials and Methods for details.

| Replicate | Treatment | Generation | n.males |
| --- | --- | --- | --- |
| 1 | C | 0 | 98 |
| 1 | N | 0 | 100 |
| 1 | P | 0 | 99 |
| 1 | Z | 0 | 100 |
| 1 | N | 1 | 95 |
| 1 | P | 1 | 100 |
| 1 | Z | 1 | 99 |
| 1 | N | 2 | 101 |
| 1 | P | 2 | 139 |
| 1 | Z | 2 | 142 |
| 1 | N | 3 | 105 |
| 1 | P | 3 | 90 |
| 1 | Z | 3 | 105 |
| 1 | N | 4 | 65 |
| 1 | P | 4 | 62 |
| 1 | Z | 4 | 67 |
| 1 | N | 5 | 97 |
| 1 | P | 5 | 100 |
| 1 | Z | 5 | 98 |
| 1 | N | 6 | 99 |
| 1 | P | 6 | 100 |
| 1 | Z | 6 | 100 |
| 1 | N | 7 | 95 |
| 1 | P | 7 | 100 |
| 1 | Z | 7 | 99 |
| 1 | N | 8 | 96 |
| 1 | P | 8 | 100 |
| 1 | Z | 8 | 99 |
| 1 | N | 9 | 90 |
| 1 | P | 9 | 100 |
| 1 | Z | 9 | 99 |
| 1 | N | 10 | 98 |
| 1 | P | 10 | 100 |
| 1 | Z | 10 | 100 |
| 1 | N | 11 | 98 |
| 1 | P | 11 | 94 |
| 1 | Z | 11 | 97 |
| 1 | C | 12 | 87 |
| 1 | N | 12 | 99 |
| 1 | P | 12 | 100 |
| 1 | Z | 12 | 99 |
| 1 | N | 13 | 98 |
| 1 | P | 13 | 100 |
| 1 | Z | 13 | 99 |
| 1 | N | 14 | 95 |
| 1 | P | 14 | 100 |
| 1 | Z | 14 | 99 |
| 1 | N | 15 | 93 |
| 1 | P | 15 | 100 |
| 1 | Z | 15 | 98 |
| 1 | N | 16 | 93 |
| 1 | P | 16 | 100 |
| 1 | Z | 16 | 98 |
| 1 | N | 17 | 127 |
| 1 | P | 17 | 132 |
| 1 | Z | 17 | 130 |
| 1 | N | 18 | 67 |
| 1 | P | 18 | 68 |
| 1 | Z | 18 | 66 |
| 1 | N | 19 | 97 |
| 1 | P | 19 | 100 |
| 1 | Z | 19 | 84 |
| 1 | N | 20 | 100 |
| 1 | P | 20 | 100 |
| 1 | Z | 20 | 100 |
| 1 | N | 21 | 99 |
| 1 | P | 21 | 100 |
| 1 | Z | 21 | 100 |
| 1 | N | 22 | 97 |
| 1 | P | 22 | 100 |
| 1 | Z | 22 | 100 |
| 1 | N | 23 | 97 |
| 1 | P | 23 | 97 |
| 1 | Z | 23 | 100 |
| 1 | N | 24 | 96 |
| 1 | P | 24 | 100 |
| 1 | Z | 24 | 97 |
| 1 | C | 25 | 70 |
| 1 | N | 25 | 72 |
| 1 | P | 25 | 122 |
| 1 | Z | 25 | 72 |
| 2 | C | 0 | 95 |
| 2 | N | 0 | 96 |
| 2 | P | 0 | 94 |
| 2 | Z | 0 | 96 |
| 2 | N | 1 | 98 |
| 2 | P | 1 | 100 |
| 2 | Z | 1 | 100 |
| 2 | N | 2 | 97 |
| 2 | P | 2 | 100 |
| 2 | Z | 2 | 100 |
| 2 | N | 3 | 95 |
| 2 | P | 3 | 100 |
| 2 | Z | 3 | 100 |
| 2 | N | 4 | 96 |
| 2 | P | 4 | 100 |
| 2 | Z | 4 | 100 |
| 2 | N | 5 | 96 |
| 2 | P | 5 | 100 |
| 2 | Z | 5 | 99 |
| 2 | N | 6 | 100 |
| 2 | P | 6 | 100 |
| 2 | Z | 6 | 100 |
| 2 | N | 7 | 96 |
| 2 | P | 7 | 100 |
| 2 | Z | 7 | 100 |
| 2 | N | 8 | 96 |
| 2 | P | 8 | 100 |
| 2 | Z | 8 | 100 |
| 2 | N | 9 | 91 |
| 2 | P | 9 | 100 |
| 2 | Z | 9 | 96 |
| 2 | N | 10 | 95 |
| 2 | P | 10 | 98 |
| 2 | Z | 10 | 99 |
| 2 | N | 11 | 98 |
| 2 | P | 11 | 100 |
| 2 | Z | 11 | 98 |
| 2 | C | 12 | 41 |
| 2 | N | 12 | 98 |
| 2 | P | 12 | 100 |
| 2 | Z | 12 | 99 |
| 2 | N | 13 | 91 |
| 2 | P | 13 | 97 |
| 2 | Z | 13 | 98 |
| 2 | N | 14 | 93 |
| 2 | P | 14 | 100 |
| 2 | Z | 14 | 100 |
| 2 | N | 15 | 97 |
| 2 | P | 15 | 100 |
| 2 | Z | 15 | 98 |
| 2 | N | 16 | 98 |
| 2 | P | 16 | 100 |
| 2 | Z | 16 | 100 |
| 2 | N | 17 | 100 |
| 2 | P | 17 | 100 |
| 2 | Z | 17 | 100 |
| 2 | N | 18 | 98 |
| 2 | P | 18 | 100 |
| 2 | Z | 18 | 100 |
| 2 | N | 19 | 99 |
| 2 | P | 19 | 100 |
| 2 | Z | 19 | 99 |
| 2 | N | 20 | 98 |
| 2 | P | 20 | 100 |
| 2 | Z | 20 | 98 |
| 2 | N | 21 | 97 |
| 2 | P | 21 | 100 |
| 2 | Z | 21 | 95 |
| 2 | N | 22 | 98 |
| 2 | P | 22 | 100 |
| 2 | Z | 22 | 99 |
| 2 | N | 23 | 99 |
| 2 | P | 23 | 100 |
| 2 | Z | 23 | 100 |
| 2 | N | 24 | 79 |
| 2 | P | 24 | 100 |
| 2 | Z | 24 | 100 |
| 2 | C | 25 | 124 |
| 2 | N | 25 | 126 |
| 2 | P | 25 | 131 |
| 2 | Z | 25 | 130 |
| 3 | C | 0 | 100 |
| 3 | N | 0 | 99 |
| 3 | P | 0 | 94 |
| 3 | Z | 0 | 99 |
| 3 | N | 1 | 96 |
| 3 | P | 1 | 98 |
| 3 | Z | 1 | 100 |
| 3 | N | 2 | 94 |
| 3 | P | 2 | 100 |
| 3 | Z | 2 | 100 |
| 3 | N | 3 | 99 |
| 3 | P | 3 | 100 |
| 3 | Z | 3 | 99 |
| 3 | N | 4 | 97 |
| 3 | P | 4 | 100 |
| 3 | Z | 4 | 99 |
| 3 | N | 5 | 94 |
| 3 | P | 5 | 100 |
| 3 | Z | 5 | 96 |
| 3 | N | 6 | 92 |
| 3 | P | 6 | 100 |
| 3 | Z | 6 | 99 |
| 3 | N | 7 | 93 |
| 3 | P | 7 | 98 |
| 3 | Z | 7 | 97 |
| 3 | N | 8 | 97 |
| 3 | P | 8 | 90 |
| 3 | Z | 8 | 98 |
| 3 | N | 9 | 96 |
| 3 | P | 9 | 100 |
| 3 | Z | 9 | 100 |
| 3 | N | 10 | 96 |
| 3 | P | 10 | 100 |
| 3 | Z | 10 | 99 |
| 3 | N | 11 | 97 |
| 3 | P | 11 | 98 |
| 3 | Z | 11 | 100 |
| 3 | N | 12 | 96 |
| 3 | P | 12 | 98 |
| 3 | Z | 12 | 99 |
| 3 | C | 13 | 95 |
| 3 | N | 13 | 91 |
| 3 | P | 13 | 94 |
| 3 | Z | 13 | 97 |
| 3 | N | 14 | 97 |
| 3 | P | 14 | 100 |
| 3 | Z | 14 | 98 |
| 3 | N | 15 | 0 |
| 3 | P | 15 | 0 |
| 3 | Z | 15 | 0 |
| 3 | N | 16 | 93 |
| 3 | P | 16 | 100 |
| 3 | Z | 16 | 99 |
| 3 | N | 17 | 96 |
| 3 | P | 17 | 100 |
| 3 | Z | 17 | 92 |
| 3 | N | 18 | 97 |
| 3 | P | 18 | 98 |
| 3 | Z | 18 | 99 |
| 3 | N | 19 | 85 |
| 3 | P | 19 | 100 |
| 3 | Z | 19 | 97 |
| 3 | N | 20 | 75 |
| 3 | P | 20 | 78 |
| 3 | Z | 20 | 92 |
| 3 | N | 21 | 86 |
| 3 | P | 21 | 96 |
| 3 | Z | 21 | 93 |
| 3 | N | 22 | 0 |
| 3 | P | 22 | 0 |
| 3 | Z | 22 | 0 |
| 3 | N | 23 | 0 |
| 3 | P | 23 | 0 |
| 3 | Z | 23 | 0 |
| 3 | N | 24 | 66 |
| 3 | P | 24 | 85 |
| 3 | Z | 24 | 73 |
| 3 | N | 25 | 0 |
| 3 | P | 25 | 0 |
| 3 | Z | 25 | 0 |
| 3 | C | 26 | 66 |
| 3 | N | 26 | 70 |
| 3 | P | 26 | 125 |
| 3 | Z | 26 | 68 |

**Table S2.**
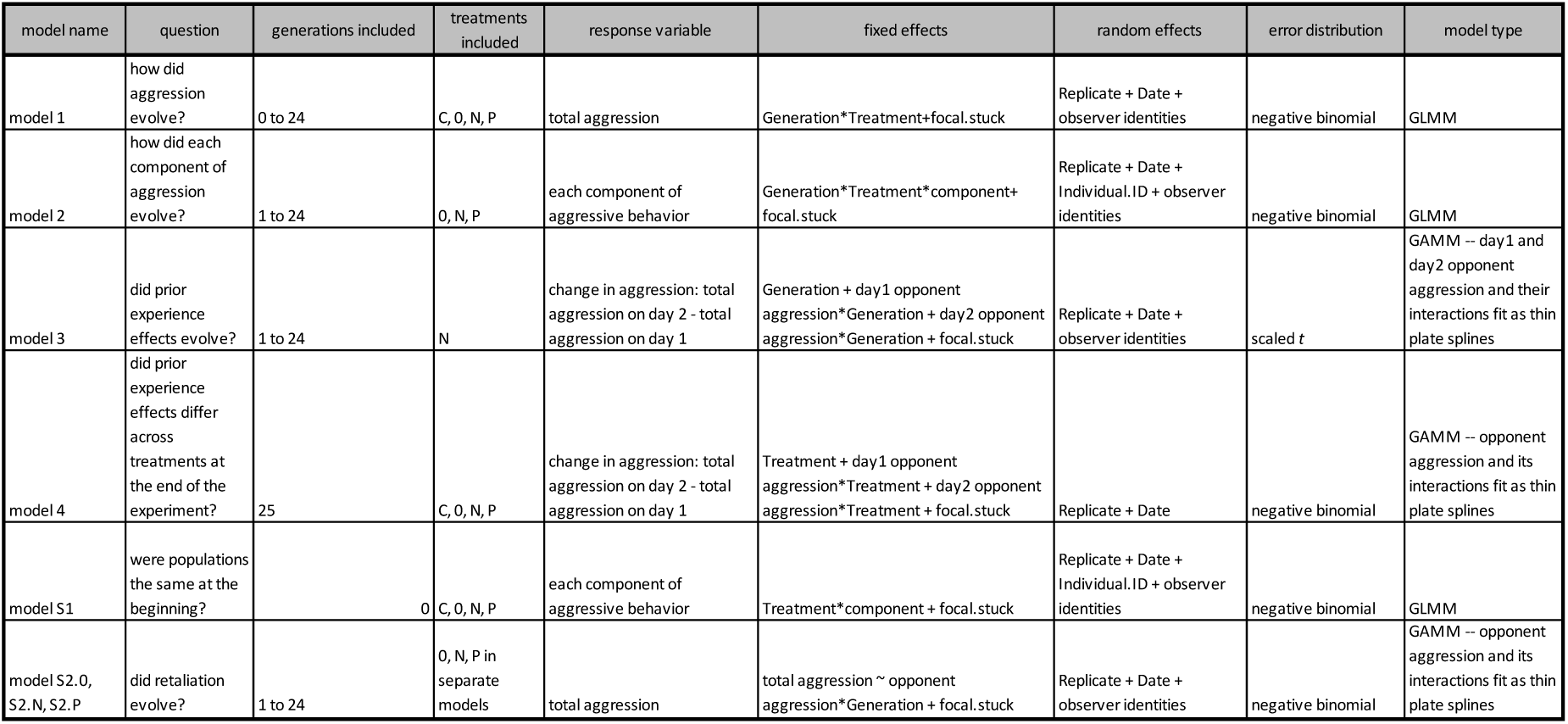
Full model structures for main and supplemental models.

| model name | question | generations included | treatments included | response variable | fixed effects | random effects | error distribution | model type |
| --- | --- | --- | --- | --- | --- | --- | --- | --- |
| model 1 | how did aggression evolve? | 0 to 24 | C, O, N, P | total aggression | Generation*Treatment+focal.stuck | Replicate + Date + observer identities | negative binomial | GLMM |
| model 2 | how did each component of aggression evolve? | 1 to 24 | O, N, P | each component of aggressive behavior | Generation*Treatment*component+focal.stuck | Replicate + Date + Individual.ID + observer identities | negative binomial | GLMM |
| model 3 | did prior experience effects evolve? | 1 to 24 | N | change in aggression: total aggression on day 2 - total aggression on day 1 | Generation + day1 opponent aggression*Generation + day2 opponent aggression*Generation + focal.stuck | Replicate + Date + observer identities | scaled t | GAMM – day1 and day2 opponent aggression and their interactions fit as thin plate splines |
| model 4 | did prior experience effects differ across treatments at the end of the experiment? | 25 | C, O, N, P | change in aggression: total aggression on day 2 - total aggression on day 1 | Treatment + day1 opponent aggression*Treatment + day2 opponent aggression*Treatment + focal.stuck | Replicate + Date | negative binomial | GAMM – opponent aggression and its interactions fit as thin plate splines |
| model S1 | were populations the same at the beginning? | 0 | C, O, N, P | each component of aggressive behavior | Treatment*component + focal.stuck | Replicate + Date + Individual.ID + observer identities | negative binomial | GLMM |
| model S2.0, S2.N, S2.P | did retaliation evolve? | 1 to 24 | O, N, P in separate models | total aggression | total aggression ~ opponent aggression*Generation + focal.stuck | Replicate + Date + observer identities | negative binomial | GAMM – opponent aggression and its interactions fit as thin plate splines |

**Table S3.** Model S1 Results.

| model S1: were populations the same at the beginning? |  |  |  |  |
| --- | --- | --- | --- | --- |
| term | parameter estimate (se) | Chisq | Df | p |
| Treatment | . | 0.1267 | 3 | 0.98846 |
| focal.stuck | -0.13 (0.078) | 2.8529 | 1 | 0.09121 |

**Table S4.** Results from Models S2.0, S2.N, and S2.P.

| model | results |  |  |  |  |
| --- | --- | --- | --- | --- | --- |
| Model 2.N | non-linear term | edf (measure of "wiggliness") | chisq | reference df | p |
|  | opponent aggression | 3.429 | 442.508 | 4.128 | < 2e-16 |
|  | Generation: opponent aggression | 1.013 | 0.122 | 1.026 | 0.739 |
| Model 2.P | non-linear term | edf (measure of "wiggliness") | chisq | reference df | p |
|  | opponent aggression | 6.298 | 785.013 | 7.115 | < 2e-16 |
|  | Generation: opponent aggression | 2.117 | 4.845 | 2.577 | 0.11528 |

| Model 2.0 | non-linear term | edf<br>(measure of "wiggliness") | chisq | reference df | p |
| --- | --- | --- | --- | --- | --- |
|  | opponent aggression | 1 | 207.259 | 1 | <2e-16 |
|  | Generation:<br>opponent aggression | 1.152 | 3.458 | 1.284 | 0.0682 |

